# Super-resolution shadow imaging reveals local remodeling of astrocytic microstructures and brain extracellular space after osmotic challenge

**DOI:** 10.1101/2021.01.05.425369

**Authors:** Misa Arizono, V.V.G. Krishna Inavalli, U. Valentin Nägerl

## Abstract

The extracellular space (ECS) plays a central role for brain physiology, shaping the time course and spread of neurochemicals, ions and nutrients that ensure proper brain homeostasis and neuronal communication. Astrocytes are the most abundant type of glia cell in the brain, whose processes densely infiltrate the brain’s parenchyma. As astrocytes are highly sensitive to changes in osmotic pressure, they are capable of exerting a potent physiological influence on the ECS.

However, little is known about the spatial distribution and temporal dynamics of the ECS that surrounds astrocytes, owing mostly to a lack of appropriate techniques to visualize the ECS in live brain tissue. Mitigating this technical limitation, we applied the recent SUper-resolution SHadow Imaging technique (SUSHI) to astrocyte-labeled organotypic hippocampal brain slices, which allowed us to concurrently image the complex morphology of astrocytes and the ECS with nanoscale resolution in a live experimental setting.

Focusing on ring-like astrocytic microstructures in the spongiform domain, we found them to enclose sizable pools of interstitial fluid and cellular structures like dendritic spines. Upon an experimental osmotic challenge, these microstructures remodeled and swelled up at the expense of the pools, effectively increasing the physical contact between astrocytic and cellular structures.

Our study reveals novel facets of the dynamic microanatomical relationships between astrocytes, neuropil and the ECS in living brain tissue, which could be of functional relevance for neuron-glia communication in a variety of (patho)physiological settings, e.g. LTP induction, epileptic seizures or acute ischemic stroke, where osmotic disturbances are known to occur.

## Introduction

Numerous studies have indicated that astrocytes can powerfully influence neuronal function by modulating the availability of neuroactive substances in the extracellular space (ECS). Astrocytes reportedly do this by rapidly removing potassium and glutamate from the ECS around synapses as well as by releasing gliotransmitters into it (1). They may also do so indirectly by molding the physical architecture of the ECS, which could effectively shape the diffusion of these substances and modulate their concentration and time course at the sites of biological action (2).

Indeed, growing evidence suggests that neuron-glia communication and brain homeostasis are regulated by astrocyte-dependent remodeling of the ECS in the healthy and diseased brain.

Whereas a tight association between astrocytic processes and dendritic spines was shown to promote glutamate uptake and gliotransmission (3–5), the withdrawal of astrocytic processes from dendritic spines after LTP reportedly boosts glutamate spillover at hippocampal synapses (6). Notably, recent work described a “glymphatic” system, where astrocytes regulate the flow of interstitial/cerebrospinal fluid, affecting the clearance of waste products from the brain (7). Moreover, astrocytic swelling and ECS shrinkage are a prominent feature of brain pathologies such as edema and epilepsy (8), and they exacerbate the disease symptoms by disturbing ion concentration gradients and fluxes and by promoting neuronal hyper-excitability via ephaptic interactions (9, 10).

While the ECS is likely to play an important role in these phenomena, it has remained a poorly characterized compartment, whose shape and dynamics around astrocytes and synapses in the neuropil remain largely unknown. Up to now, the ECS has mostly been studied by iontophoretic or fluorescence techniques that quantify the radial spread of diffusible probes released from a point source (11). While these techniques provide precise measurements of bulk biophysical parameters such as volume fraction and tortuosity, they do so over relatively large chunks of brain tissue (>100 *µ*m) and cannot reveal the spatial structure of the ECS. Electron microscopy (EM) offers enough spatial resolution, but it requires tissue fixation, which precludes following live events through time and may introduce artifacts by distorting the tissue and depleting the ECS depending on the fixation protocol (12).

We sidestepped these technical limitations by using the recent SUSHI technique (SUper-resolution SHadow Imaging; (13)), which can visualize the anatomical organization of the ECS and, by implication, all cellular structures with very high resolution and contrast in living brain tissue, because the ECS is basically the negative imprint of the brain cells. It relies on labeling of the interstitial fluid in the ECS with a small but membrane-impermeant fluorescent dye, which casts all cellular structures, including their finest processes, as a sharp shadow under a 3D-STED microscope. Combining SUSHI with positive fluorescence labeling of astrocytes, we could tease apart the ECS and astrocytes from all other cellular structures, allowing us to examine their dynamic anatomical relationships in living organotypic brain slices.

Focusing on recently described ring-like astrocytic microstructures (14), we observed that they frequently enclosed pools of interstitial fluid in the ECS and synaptic structures like dendritic spines. Upon osmotic challenge, which caused the astrocytes to swell, the aperture of the rings constricted at the expense of the ECS pools, effectively increasing their contact with the cellular structures inside.

## Results

### Imaging strategy to reveal ECS and astrocyte fine structure

To resolve the fine structure of astrocytes and ECS in organotypic hippocampal brain slices, we employed our recent SUSHI technique based on a custom-built 3D STED microscope (13), which provides a volume resolution of under 1 attoliter (60 nm in x,y and 200 nm in z). The slices were prepared from crossbred transgenic mice (Cre/ERT2 and CAG-Floxed ZsGreen), where all astrocytes expressed the fluorescent protein ZsGreen upon hydrotamoxifen treatment (14).

For concurrent visualization of astrocytes and the ECS, we added the fluorescent dye ATTO514 to the ACSF perfusate. We applied linear unmixing to the spectrally detected fluorescence signals, which reduced their crosstalk to under 10% (see Methods for details and (15). To facilitate the perception of contrast and forms in the images of the densely labeled tissue, we inverted the ATTO514 signal, which more readily reveals the myriad anatomical relationships between ECS, astrocytes and the other cellular structures in the tissue (Fig.1A). We focused our analysis on the stratum radiatum of the CA1 area of the hippocampus, where neurites and astroglial processes strongly intermingle, forming ‘tripartite’ synapses (Fig.1B), which have been under intense investigation by neurophysiologists (16, 17).

### Astrocytic microstructures enclose ECS pools and cellular structures

In the dually labeled tissue of astrocytes and ECS (Fig. 2A), it was possible to discern cellular structures like dendrites, spines and astrocytic processes and to distinguish them from the surrounding ECS (Fig. 2B).

**Fig 1.**
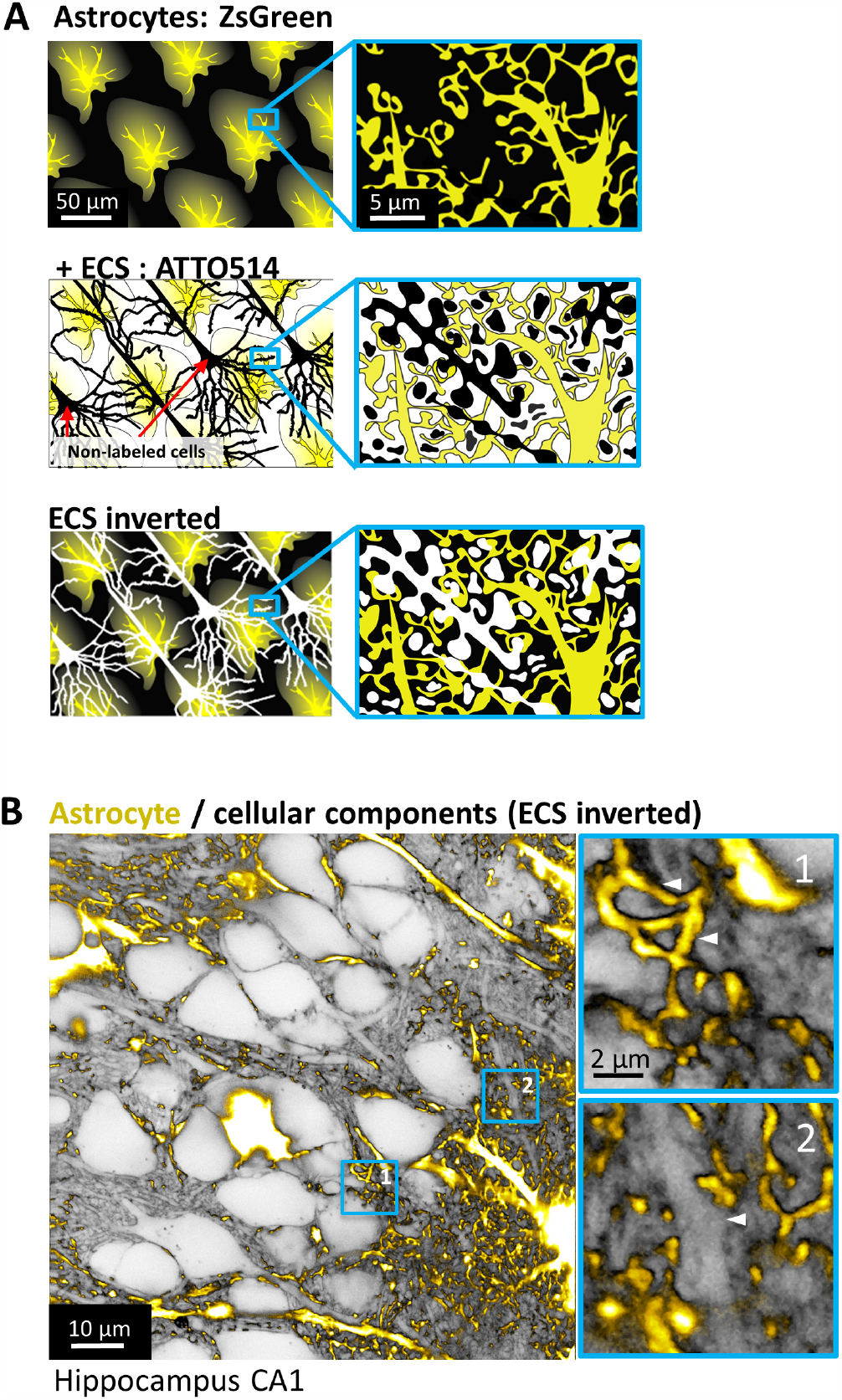
Imaging strategy to reveal ECS and astrocyte fine structure. **A**. Schematic of astrocyte and ECS imaging using STED microscopy. We used organotypic hippocampal slices where astrocytes are labeled with fluorescent protein ZsGreen (top). By adding fluorescent dye (ATTO514) in the extracellular solution, we can visualize the extracellular space and astrocytes (middle). Inverting the signal reveals the relationship between astrocytes and all the cellular structures in the tissue (bottom). **B**. A 3D STED image revealing astrocytic processes infiltrating the neuropil visualized by inverting the ECS image. Zoom-in images show astrocytic processes enwrapping round cellular structures (1, white triangles) and contacting a putative dendrite (2, white triangle).

**Fig 2.**
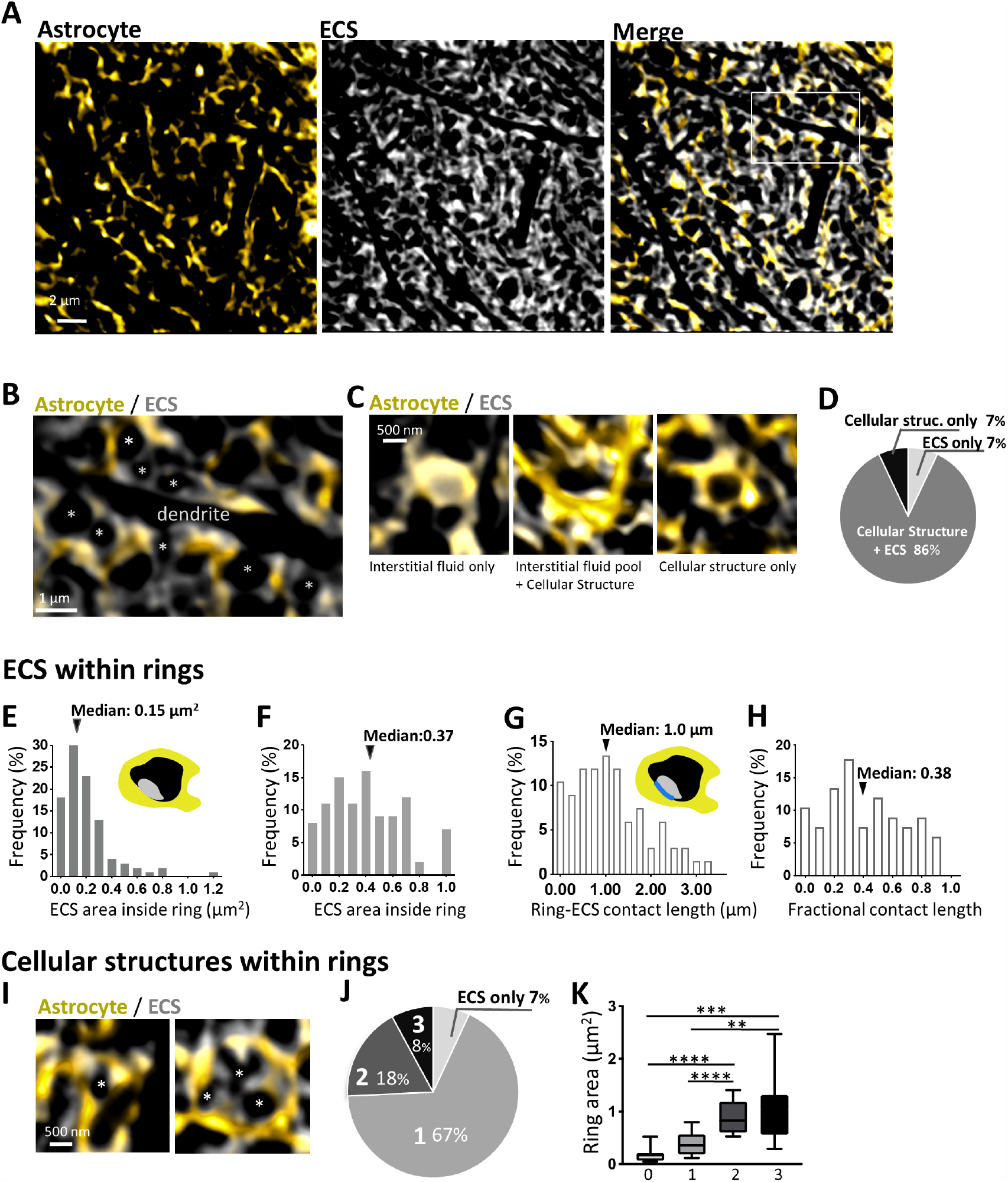
Astrocytic microstructures enclose interstitial pools and cellular structures. **A**. STED images of reticular structure of astrocytes and ECS visualized by combining positive labeling of astrocytes and staining of ECS. **B**. A STED image of astrocyte and ECS showing a negative imprint of a putative dendrite, spines (*) (black), perisynaptic astrocytic processes (yellow) and abundant ECS (grey). **C**. Images of rings with only interstitial fluid (left), a pool of interstitial fluid (middle) and without any detectable interstitial fluid (right). **D**. Percentage of rings depending on the encircled structure presented in C. **E**. Distribution of ECS area within the rings. **F**. Distribution of percentage of ECS area within the rings. **G**. Distribution of inner perimeter of the rings in direct contact with the interstitial fluid pool. **H**. Distribution of percentage of inner perimeter of the ring in direct contact with the interstitial fluid pool. **I**. Example of rings enwrapping cellular structures with simple round geometry (left), and elaborate geometry which may reflect multiple cellular structures (right). **J**. Percentage of rings enwrapping 0, 1, 2, and 3 distinguishable negative imprints. **K**. Comparison of area inside the ring which are enwrapping 0 (n = 7) 1 (n = 67), 2(n = 18), and 3(n = 8) negative imprints. Larger loops enwrap multiple cellular components (5 slices; Kruskal-Wallis test, **p < 0.01, ***p <0.001, ****p < 0.0001).

We focused our attention on the ring-like astrocytic microstructures that we recently identified as a ubiquitous feature of protoplasmic astrocytes in the mouse brain (14). They are circular arrangements of shafts and nodes that appear to reconnect and form closed rings that infiltrate the neuropil as a close-knit meshwork.

We observed that the rings enclosed a variable amount of (non-astrocytic) cellular structures, which oftentimes appeared to be dendritic spines, and pools of ECS filled with interstitial fluid (Fig. 2C). Indeed, these cellular structures are comparable to dendritic spine heads in size (Fig. S1A), and by dual labeling of astrocytes and neurons, we confirmed that dendritic spines can be seen inside the rings (Fig. S1B). Most of the time (86 %) the astrocytic rings enclosed both cellular structures and interstitial fluid pool at the same time, but sometimes they appeared to enclose only cellular structures (7 %) or only interstitial fluid (7 %) (Fig. 2D).

The median area of the ECS pools enclosed by the rings was 0.15 µm2 (Fig. 2E), corresponding to 37 % of the ring area (Fig. 2F), while the median length of the contact zone between ring and ECS pool was around 1 *µ* (Fig. 2G) corresponding to 38 % of the ring perimeter (Fig. 2H). The analysis shows that the ECS takes up a sizable portion and not just a thin lining along inner surface of the astrocytic rings. Moreover, rings were smaller when they only enclosed ECS and most rings contained only a single cellular structure of simple shape, but larger rings were seen to enclose also multiple or complex shaped structures (Fig. 2I-K).

### Astrocytic microstructures remodel after osmotic challenge, depleting local ECS

Next, we investigated how osmotic challenges might impact this arrangement of astrocytic microstructures, cellular structures and ECS, given that astrocytes are known to be highly sensitive to osmotic disturbances induced by strong neuronal activity or ischemic conditions (18, 19).

Benefitting from the live-cell imaging approach, we imaged the labeled slices before and during the application of hypoosmotic conditions (HOC). The osmotic challenge consisted of reducing the osmolarity (i.e. the total number of solute particles per liter of solution) of the standard-formula perfusate (ACSF), from 300 mOsmol to 200 mOsmol for a duration of 30 minutes (Fig. 3A). This is considered a relatively severe form of osmotic stress (18) and similar condition (−30 % mOsm) was reported to cause sustained astrocytic swelling in vivo (20).

**Fig 3.**
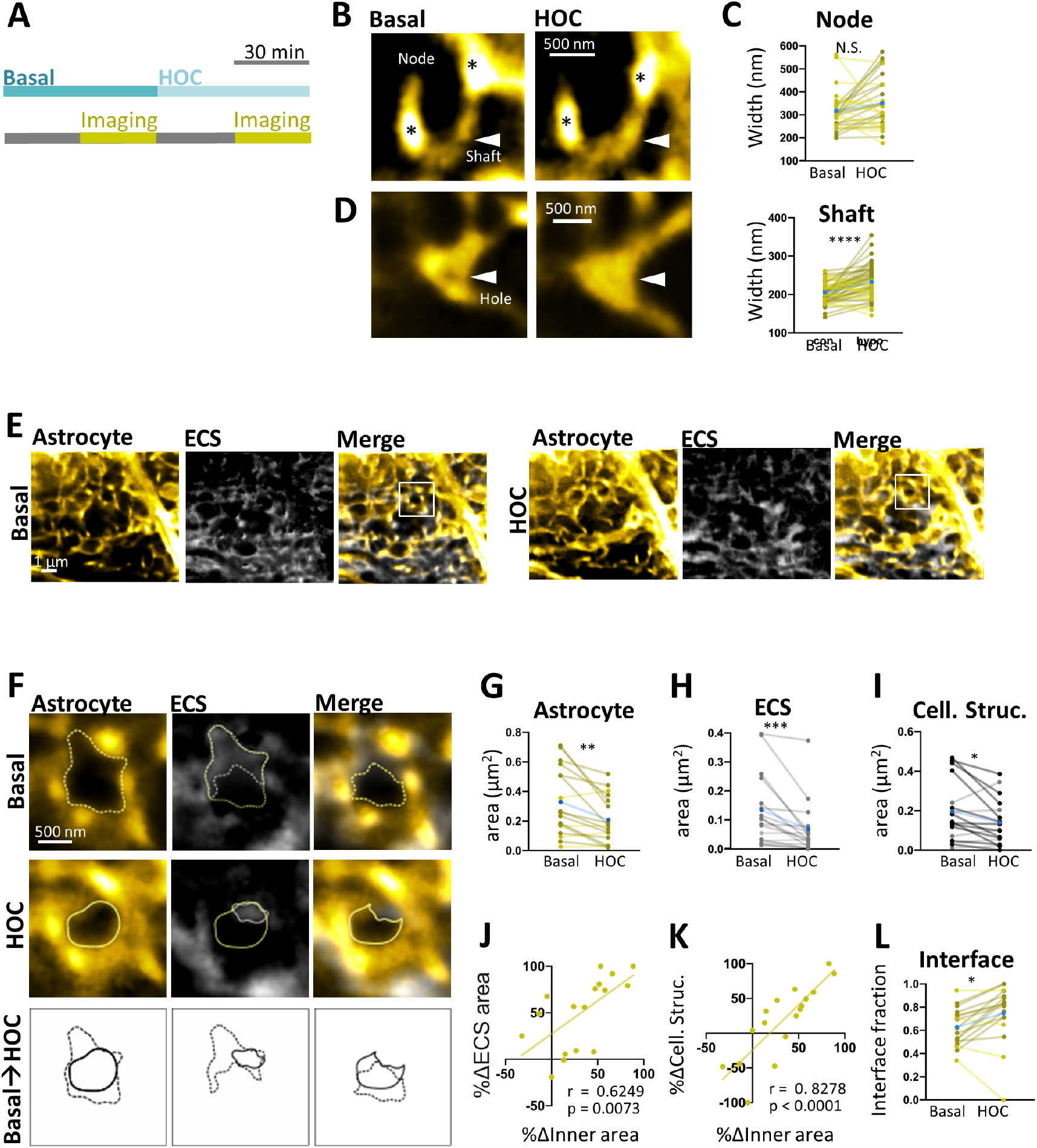
Astrocytic microstructures remodel after osmotic challenge, depleting local ECS. **A**. Time line of the experiment. **B**. STED images of astrocytic processes before (basal) and after hypo-osmotic condition (HOC). Shafts (triangle) connecting nodes (*) selectively swelled. **C**. Top: Comparison of astrocytic node width before (basal) and after HOC (n = 30 from 4 slices; Wilcoxon test, N.S. Not Significant). Bottom: Comparison of astrocytic shaft width before (basal) and after HOC (n = 52 from 4 slices; Wilcoxon test, ****p<0.0001). **D**. STED images of astrocytic ring before (basal) and after HOC. The hole in the center of the astrocytic ring (white triangle) appears to close in HOC. **E**. STED images of reticular structure of an astrocyte and ECS, before (basal, left) and after (HOC, right) HOC. **F**. Left: Magnified image of E showing a representative ring (left), encircled ECS (middle), and negative imprint of a cellular structure (right). The bottom panels show the changes before (dotted line) and after (solid line) hypo-osmotic stress. **G-I**. Total ring inner area (G), encircled ECS area (H), encircled cellular structure area (I) before (basal) and after HOC (n = 17 from 3 slices; Wilcoxon test, *p < 0.05, **p < 0.01, ***p < 0.001,). **J**. Correlation between fraction change of the ring inner area and fraction change of encircled ECS (n = 17 from 3 slices; Spearman r = 0.6249, p = 0.0073). **K**. Correlation between fraction change of the ring inner area and fraction change of encircled cellular structure (n = 17 from 3 slices; Spearman r = 0.8278, p < 0.0001). **L**. Fraction of inner perimeter of the loop in contact with cellular structure before and after HOC (n=17 from 3 slices; Wilcoxon test, *p < 0.05). For paired analysis (C, G, H, I, L), blue points indicate the mean values, those that decreased or increased are shown in different shades of color to demonstrate the heterogenous nature of the response.

The astrocytic microstructures reacted to the osmotic challenge by remodeling their shape and volume. The morphological changes were heterogeneous in magnitude and direction as well as in the type of structures that were impacted. The astrocytic nodes showed a mixed and modest response, with about 43 % of them undergoing shrinkage while 57 % showed swelling (Fig. 3B,C). By contrast, the astrocytic shafts exhibited a more consistent response. Their diameters increased from 204 nm to 231 nm (median) following the osmotic challenge, which was statistically highly significant (mean ± sem, basal: 204.1 ± 3.9; HOC: 231.5 ± 6.0; p < 0.0001 Wilcoxon matched-pairs signed rank text; N = 51; Fig. 3B,C). This difference was absent under control conditions where hypo-osmotic stress was not applied (Fig. S2A,B), showing that the swelling was not caused by STED imaging or the long incubation time in the chamber.

Notably, in 21 % of the cases, the ring-like organization of the astrocytic processes disappeared after the osmotic challenge, suggesting a closure of the holes within the rings (Fig. 3D).

By co-labeling the ECS, we could also observe how the ECS was affected by the astrocytic remodeling (Fig. 3E). We observed that the inner area of the rings decreased (from 0.33± 0.056 *µ*m^2^ to 0.21 ± 0.039 *µ*m^2^ (mean ± sem), Wilcoxon matched-pairs signed rank text; p = 0.0010, N = 17; Fig. 3F,G), consistent with the swelling of the astrocytes, as they delineate the perimeter of this area. At the same time, the areas occupied by the ECS and cellular structures inside the rings decreased substantially (ECS, from 0.13 ± 0.030 *µ*m^2^ to 0.066 ± 0.022 *µ*m^2^ (mean ± sem), Wilcoxon matched-pairs signed rank text; p = 0.0001, N = 17; Fig. 3F, H; cellular structure, from 0.20 ± 0.038 *µ*m^2^ to 0.14 ± 0.029 *µ*m^2^ (mean ± sem), Wilcoxon matched-pairs signed rank text; p = 0.023, N = 17; Fig. 3F,I).

The decrease in ring area was highly correlated with the decrease in ECS (Fig. 3J) and cellular areas inside the ring (Fig. 3K), mutually corroborating the independent measurements. We observed that the zone of contact between the inner side of the astrocytic rings and the cellular structures increased during osmotic challenge, which might affect the ability of these structures to interact physically or exchange molecular signals at short range (interface fraction, from 0.63± 0.040 to 0.75 ± 0.061 (mean ± sem), Wilcoxon matched-pairs signed rank text; p = 0.011, N = 17; Fig. 3L).

## Discussion

We employed 3D super-resolution microscopy in two colors for volumetric imaging of astrocytes and ECS in living brain slices to reveal their dynamic anatomical relationships induced by an osmotic challenge. The inverse labeling strategy of SUSHI also revealed the surrounding anatomical landscape, making it possible to identify and monitor the neighboring cellular structures alongside the positively labeled astrocytes.

Much of what we know about astrocytic dynamics and swelling (20–22) has come from studies using light microscopy. However, these studies only reported on the cell bodies and major branches of the astrocytes because of the diffraction limit of light microscopy, which makes it impossible to resolve the spongiform domain, where their highly ramified processes contact synapses in the neuropil. Quantitative biophysical techniques like TMA^+^ iontophoresis have been used to measure changes in the global volume fraction of the ECS as a surrogate for astrocytic swelling (23), but they can only indicate net changes in overall cellular volume and are insensitive to shape changes.

By contrast, our new approach can directly track even the finest individual astrocytic structures over time and read out changes in their local volume and shape with very high spatial resolution, while making it possible to monitor the ECS, and thus, albeit indirectly, also all other cellular structures at the same time.

We focused our attention on ring-like astrocytic microstructures, which are formed by ‘shafts’ and ‘nodes’ of circularly connecting astrocytic processes we described recently (14). The rings typically enclosed cellular structures (such as dendritic spines) and contained sizable pools of interstitial fluid. Upon osmotic challenge, the astrocytic shafts grew wider, reducing or even closing the aperture of the rings. At the same time, the enclosed ECS and cellular structures shrank or were pushed out of the rings. The astrocytic microstructures tended to take on less complex shapes during osmotic stress, which could be a way to accommodate a larger cell volume without having to add new cell membrane.

The astrocytic remodeling we observed is likely a direct biophysical consequence of the change in osmotic pressure, which causes a net water movement into the astrocytes, causing them to swell up at the expense of the ECS pools (reviewed in (24)). Interestingly, the nodes remained largely unaffected, suggesting their membrane or cytoskeletal organization have special properties that provide resistance to osmotically induced structural remodeling.

The effects we have observed could have several implications for neuronal function. The local depletion of the ECS may increase the concentration levels of extracellular K^+^ and glutamate, and thereby promote neuronal excitability and synaptic transmission (18). The astrocytic swelling and the resultant morphological ‘de-complexification’ may alter the spatial organization of the tripartite synapse and thereby affect the spread of glutamate and its uptake around synapses. The widening of astrocytic shafts may also facilitate the spread of Ca^2+^ signals inside the astrocytes (14), modulating the release of ‘gliotransmitters’ like glutamate. Finally, the swelling of the astrocytic rings may also generate mechanical forces that physically displace adjacent cellular structures and activate mechanosensitive receptors in their membranes (25).

Our study may be relevant for understanding brain damage such as traumatic brain injury and stroke (26), providing a more refined picture of how astrocytic swelling manifests itself within a shifting anatomical landscape of nanoscale dimensions. A variety of molecules such as gap junction proteins (Connexins 30 and 43), Aquaporins, TRP channels, VRAC channels have been implicated in mediating structural remodeling of astrocytes, however their exact contributions remain unknown (8), which is partly due to the technical difficulties in resolving the underlying anatomical structures. Applying our novel super-resolution / SUSHI imaging approach to the intact mouse brain in vivo will open up unprecedented experimental opportunities to study the complex phenomenology, molecular mechanisms and functional consequences of astrocytic swelling in a variety of pathophysiologically relevant settings and disease models.

## Materials and Methods

### Organotypic hippocampal slice cultures

Organotypic hippocampal slices (27) were dissected from 5–7 days old mice, obtained from cross-breeding GFAP-cre (MGI:4418665) and Ai6 (JAX: SN007906) mice (13), and were cultured 5–8 week in a roller drum at 35 °C. ZsGreen expression was induced by 1µM 4-Hydroxytamoxifen (Sigma-Aldrich) treatment once per week for 5 weeks (14). Experimental procedures were in accordance with the European Union and CNRS UMR5297 institutional guidelines for the care and use of laboratory animals (Council directive 2010/63/EU).

### Viral infection of organotypic slices

Microinjectionof Sindbis-Citrine virus into the hippocampal slice for neuronal labeling was performed using a glass pipette connected to Picospritzer (Parker Hannifin). In brief, the virus was injected via a pipette positioned into the CA1 area of the slice by brief pressure pulses (40ms; 15 psi) one day prior to the experiment (14).

### Hypo-osmotic challenge and ECS staining

Baseline experiments were performed in artificial cerebrospinal fluid (ACSF) containing 119 mM NaCl, 2.5 mM KCl, 1.3 mM MgCl_2_, 2.5 mM CaCl_2_, 26 mM NaHCO_3_, 1 mM NaH_2_PO_4_, 20 mM D-glucose, 1 mM Trolox; 300 mOsm; pH 7.3. Hypoosmotic stress was applied by perfusing ACSF with reduced NaCl concentration (119 mM to 69 mM, 300 mOsm to 200 mOsm). Perfusion rate was 2 ml/min and the temperature was 32 °C. For ECS staining, ATTO-514 (final concentration 40 µM) was pipetted directly into the chamber while pausing the perfusion.

### 3D STED microscopy

We used a home-built 3D STED microscope constructed around an inverted microscope body (Leica DMI6000 CS) with a 1.3 NA glycerol immersion objective (HC PL APO 63X, Leica Microsystems, Germany) equipped with a correction collar to compensate spherical aberrations and thus to allow imaging deeper inside tissue as described in detail previously (28). The microscope uses pulsed excitation at 488 nm (PDL 800-D, PicoQuant) and pulsed quenching at 594 nm. The quenching wavelength is obtained from an optical parametric oscillator pumped at 732 nm by a Ti-sapphire femtosecond laser (MaiTai, SpectraPhysics, US). The 3D resolution enhancement was achieved by splitting the STED beam into two beams; imposing one to a 2*π* delay vortex phase mask for lateral resolution and the other to an annular π delay phase mask for axial resolution, and subsequently recombining them, as described in detail previously (13). The combination of ZsGreen and ATTO 514 signals were spectrally separated using a 514 nm long-pass dichroic mirror and detected with two avalanche photodiodes (15).

### Image acquisition

STED z-stack images were typically acquired with the imaging conditions (1) 20 *×*20 *×*2*µm*^3^, pixel size 19.53 nm, z plane 80 nm × 25 planes for single z-stacks, and (2) 10 *×*10 *×*0.5*µm*^3^, pixel size 19.53 nm, z plane 80 nm *×*5 planes for paired image acquisitions in basal and hypoosmotic conditions. Image acquisition was controlled by the Imspector software.

### Image processing and analysis

All STED images were deconvolved by Huygens Professional (SVI) software. ZsGreen (astrocyte) and ATTO 514 (ECS) or Citrine (neuronal labeling) were spectrally unmixed with ImageJ software (NIH) (15). To measure astrocytic process width, we used the ImageJ plugin-SpineJ (29). For parameters regarding the rings, the analysis was performed by manual tracing of the structures and subsequently calculating the area or perimeter using ImageJ software (NIH).

## ACKNOWLEDGEMENTS

The study was supported by research grants from the Agence Nationale de la Recherche (ANR-12-BSV4-0014 and ANR-17-CE16-0002) and Fondation pour la Recherche Médicale (DEQ20160334901). MA was supported by a postdoctoral fellowship from JSPS (Japan). We thank J. Angibaud for preparing organotypic slices. We thank J. Badaut and all members of the Nägerl team for comments on the manuscript.

---

**Fig S1.**
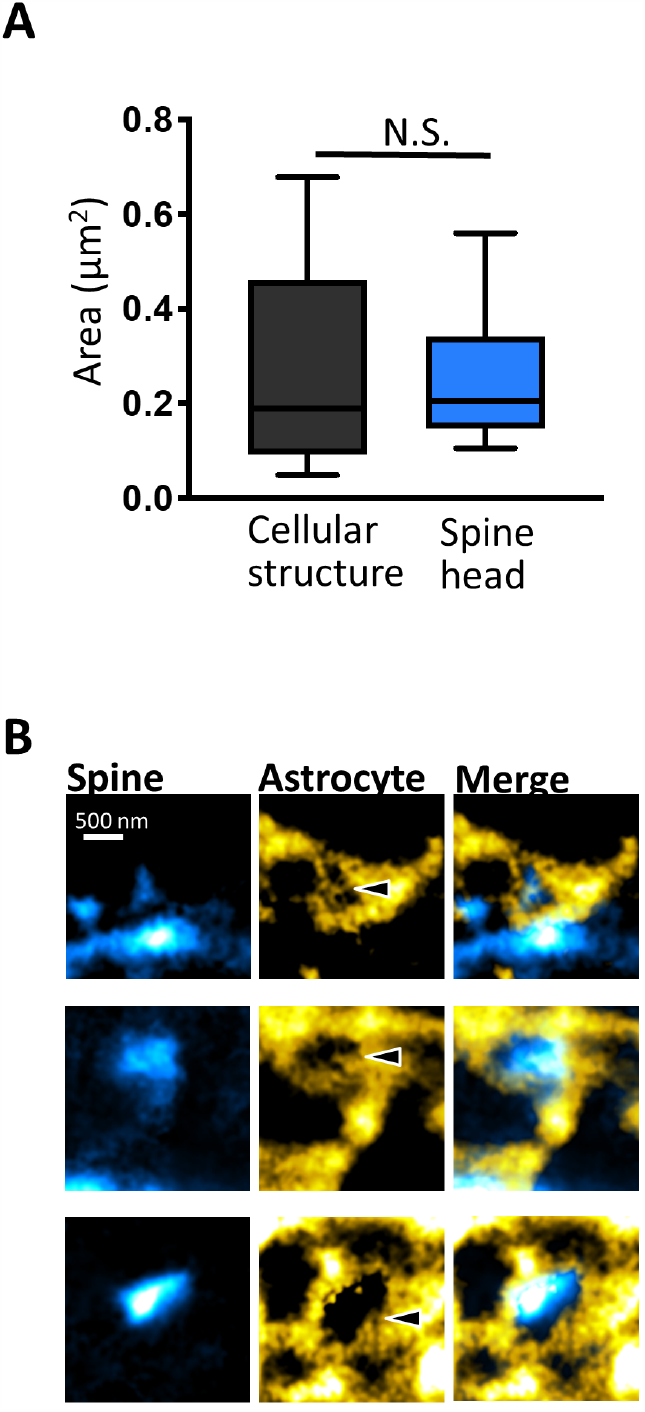
Spines inside astrocytic rings. **A**. Size comparison between encircled cellular structure and positively labeled spine head (n = 68 from 5 slices, n = 104 from 21 slices; Mann-Whitney test, p = 0.35, N.S. Not Significant). **B**. Dual labeling of dendrites (cyan) and astrocytes (yellow) reveals dendritic spines can be enclosed by astrocytic rings. Note that only a subset of neurons is labeled in this sample.

**Fig S2.**
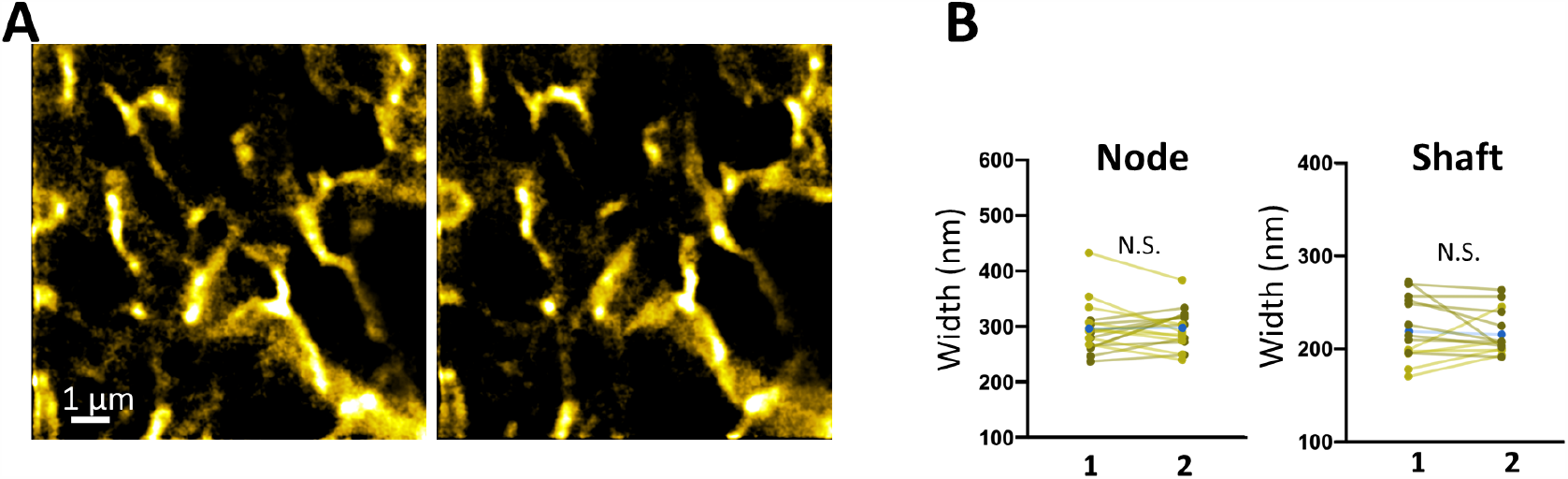
Swelling is absent under control conditions. **A**. STED images of astrocytes in control conditions without hypo-osmotic stress. Left: First STED acquisition Right: Second STED acquisition taken after 1 hour. **B**. Left: Comparison of astrocytic node width between first (1) and second (2) image acquisition (n = 18 from 3 slices; Wilcoxon test, p = 0.87, N.S. Not Significant). Right: Comparison of astrocytic shaft width between first (1) and second (2) image acquisition (n = 14 from 3 slices; Wilcoxon test, p = 0.63, N.S. Not Significant).

## Notes

### Competing Interest Statement

The authors have declared no competing interest.

